# Age-related increases in reaction time result from slower preparation, not delayed initiation

**DOI:** 10.1101/2021.06.12.448183

**Authors:** Robert M Hardwick, Alexander D. Forrence, Maria Gabriela Costello, Kathy Zackowski, Adrian M Haith

## Abstract

Recent work indicates that healthy younger adults can prepare accurate responses faster than their voluntary reaction times indicate, leaving a seemingly unnecessary delay of 80-100ms before responding. Here we examined how the preparation of movements, initiation of movements, and the delay between them are affected by ageing. Participants made planar reaching movements in two conditions. The ‘Free Reaction Time’ condition assessed the voluntary reaction times with which participants responded to the appearance of a stimulus. The ‘Forced Reaction Time’ condition assessed the minimum time actually needed to prepare accurate movements by controlling the time allowed for movement preparation. The time taken to both initiate movements in the Free Reaction Time and to prepare movements in the Forced Response condition increased with age. Notably, the time required to prepare accurate movements was significantly shorter than participants’ self-selected initiation times; however, the delay between movement preparation and initiation remained consistent across the lifespan (~90ms). These results indicate that the slower reaction times of healthy older adults are not due to an increased hesitancy to respond, but can instead be attributed to changes in their ability to process stimuli and prepare movements accordingly, consistent with age-related changes in brain structure and function.

## Introduction

Adult human reaction times in response to simple tasks slow with age at a rate of 2-6ms per decade (Fozard et al. 1994; Gottsdanker 1982; Woods et al. 2015). More complex tasks are associated with greater reaction time differences between healthy young and old participants (Woods et al. 2015). These increases in response times have been attributed to changes in both the physical capabilities and the self-selected behaviors of older adults. Age-related changes in brain physiology are associated with reductions in the speed of information processing (Seidler et al. 2010). Compared to younger adults, older individuals have reduced grey matter volumes (Giorgio et al. 2010), reductions in white matter integrity (Stadlbauer et al. 2008), and recruit additional neural resources when completing tasks (Heuninckx et al. 2008), all of which could contribute to slower sensorimotor processing times. A second factor that may contribute to this decline comes from research suggesting that older adults take a more cautious approach when performing tasks (Dully et al. 2018). For tasks in which performance is governed by a speed-accuracy trade-off (Fitts 1954), younger adults appear to balance speed and accuracy in a way that achieves a high rate of correct responses, while older adults reportedly focus on minimizing errors at the cost of being slower (Salthouse 1979; Smith and Brewer 1995; Starns and Ratcliff 2010). It is unclear which of these explanations – slower processing or greater cautiousness – is primarily responsible for the general increase in reaction times with ageing.

Cautiousness to respond (i.e. focusing on accuracy over speed) appears to occur even in tasks that one might expect to be highly reactive, such as reaching to a visual target. We have recently shown that healthy younger adults can detect a target location and prepare an accurate movement in as little as 150ms, but introduce a delay of 80-100ms before voluntarily initiating a response (Haith et al. 2016), seemingly to avoid committing errors in which responses were initiated before they had been prepared. Here our goal was to quantify the effects of aging on movement preparation, movement initiation, and the relationship between them. We hypothesized that if healthy older adults delay their actions in order to favor accuracy, the delay between the minimum time required to prepare movements and the time at which they are voluntarily initiated may increase with age.

In the present study we therefore examined the extent to which the slower reaction times of healthy older individuals are due to a slowing of their ability to process perceptual information and prepare appropriate movements (i.e. due to an overall reduction in processing speed), and/or an increase in the delay between when their movements are prepared and initiated (e.g. favoring accuracy over speed to avoid the risk of making an error). Participants completed a planar reaching task, and their reaction times were measured in two different conditions. The ‘Free Reaction Time’ condition (equivalent to standard “choice reaction time” testing), assessed the time at which participants would voluntarily initiate movements in response to the appearance of a target. The ‘Forced Reaction Time’ condition, based on an established psychophysics paradigm (Ghez et al. 1997; Haith et al. 2015a, 2015b, 2016; Hardwick et al. 2019), forced participants to respond at lower-than-normal reaction times, allowing us to determine the amount of time they needed to prepare accurate responses. Our results indicate that the time participants required to both initiate and prepare responses increased with age; however, the delay between preparation and initiation of movements remained invariant at around 90ms. These results indicate that the slower reaction times of healthy older adults observed in this task were not due to an increased hesitancy to respond, but can instead be wholly attributed to declines in the ability to process stimuli and prepare accurate movements.

## Methods

54 human participants aged between 21-80 completed the study (see Table 1 for summary data). Previous research indicates typical correlations between age and reaction time in the range of r=0.46 to r=0.51 (Bugg et al. 2006; Woods et al. 2015). Power analysis based on the more conservative r=0.46, with 80% power and a two-tailed alpha of 0.05 indicated that a sample of 35 participants would be sufficient to detect effects in the present study (based on power analysis calculations from Hulley 2007). All participants had no known neurological disorders and had normal cognition (a score of ≥26 on the Montreal Cognitive Assessment (Nasreddine et al. 2005). All participants provided written informed consent prior to their participation, and all procedures were approved by the Johns Hopkins University School of Medicine Institutional Review Board.

**Table 1:** Test population summary data

|  | Age | Sex | Hand | Initiation Time<br>(Mean±SD, ms) | Mean PT (ms) | Difference<br>(IT - PT) | Accuracy<br>(Free RT, %) |
| --- | --- | --- | --- | --- | --- | --- | --- |
|  | 21 | F | R | 355±43 | 264 | 91 | 95.3 |
|  | 21 | F | R | 399±36 | 280 | 119 | 99.0 |
|  | 21 | F | R | 387±27 | 283 | 104 | 100.0 |
|  | 21 | F | L | 364±38 | 292 | 72 | 99.7 |
|  | 21 | F | R | 420±47 | 304 | 116 | 100.0 |
|  | 23 | M | R | 380±32 | 305 | 75 | 99.2 |
|  | 24 | M | R | 368±36 | 289 | 79 | 98.7 |
|  | 25 | F | R | 346±49 | 238 | 108 | 99.3 |
|  | 28 | M | R | 370±29 | 268 | 102 | 100.0 |
|  | 28 | F | R | 387±31 | 275 | 112 | 96.9 |
|  | 29 | M | R | 344±27 | 253 | 91 | 99.7 |
|  | 29 | F | L | 328±55 | 271 | 57 | 99.0 |
|  | 29 | M | R | 403±29 | 271 | 132 | 99.5 |
|  | 30 | F | R | 329±24 | 255 | 74 | 100.0 |
|  | 31 | F | R | 398±31 | 294 | 104 | 99.5 |
|  | 34 | M | R | 354±28 | 294 | 60 | 99.5 |
|  | 35 | M | R | 339±20 | 265 | 74 | 99.5 |
|  | 36 | F | R | 400±82 | 286 | 114 | 100.0 |
|  | 36 | F | R | 404±53 | 327 | 77 | 98.2 |
|  | 37 | F | R | 435±38 | 303 | 132 | 98.4 |
|  | 38 | F | R | 391±37 | 315 | 76 | 99.7 |
|  | 40 | F | L | 395±36 | 281 | 114 | 99.2 |
|  | 40 | M | R | 411±59 | 302 | 109 | 97.9 |
|  | 42 | F | R | 411±39 | 293 | 118 | 100.0 |
|  | 43 | F | R | 352±41 | 291 | 61 | 100.0 |
|  | 44 | F | R | 370±41 | 309 | 61 | 98.4 |
|  | 45 | F | R | 429±31 | 328 | 101 | 99.0 |
|  | 45 | M | R | 480±17 | 339 | 141 | 99.5 |
|  | 47 | F | R | 380±48 | 300 | 80 | 98.4 |
|  | 50 | F | R | 404±42 | 325 | 79 | 96.4 |
|  | 55 | F | R | 444±35 | 288 | 156 | 98.7 |
|  | 56 | F | R | 370±42 | 270 | 100 | 99.7 |
|  | 56 | F | R | 327±37 | 288 | 39 | 100.0 |
|  | 56 | M | R | 421±42 | 338 | 83 | 99.7 |
|  | 57 | F | R | 394±77 | 284 | 110 | 96.5 |
|  | 57 | F | R | 372±39 | 289 | 83 | 99.0 |
|  | 58 | M | R | 411±34 | 309 | 102 | 96.9 |
|  | 59 | F | R | 429±40 | 278 | 151 | 99.5 |
|  | 59 | F | R | 379±35 | 321 | 58 | 99.0 |
|  | 59 | M | R | 362±69 | 323 | 39 | 100.0 |
|  | 61 | F | R | 378±30 | 299 | 79 | 94.5 |
|  | 62 | F | L | 431±29 | 298 | 133 | 99.5 |
|  | 62 | M | R | 361±35 | 305 | 56 | 99.2 |
|  | 63 | M | R | 439±35 | 299 | 140 | 98.7 |
|  | 66 | F | R | 351±44 | 228 | 123 | 100.0 |
|  | 68 | M | R | 384±34 | 282 | 102 | 99.5 |
|  | 70 | F | L | 468±33 | 382 | 86 | 100.0 |
|  | 71 | F | R | 379±41 | 294 | 85 | 100.0 |
|  | 72 | M | R | 381±45 | 294 | 87 | 98.7 |
|  | 72 | M | R | 386±38 | 314 | 72 | 99.0 |
|  | 72 | F | R | 383±34 | 315 | 68 | 99.0 |
|  | 72 | M | L | 397±42 | 326 | 71 | 96.4 |
|  | 76 | F | R | 400±54 | 301 | 99 | 97.9 |
|  | 80 | M | R | 453±32 | 325 | 128 | 100.0 |
| Summary | Mean±SD | Count | Count | Mean±SD | Mean±SD | Mean±SD | Mean±SD |
|  | 46.9±17.6 | 35F, 19M | 48R, 6L | 390±35ms | 295±26ms | 94±28ms | 98.9±1.3% |

### Apparatus

Participants sat at a glass-surfaced table with their dominant arm supported by an air sled, allowing frictionless 2D movements in the horizontal plane (see Figure 1). A monitor and mirror setup allowed presentation of visual targets in the same plane as the arm. Hand position was tracked at 130Hz using a Flock of Birds motion tracking system (Ascension Technologies).

**Figure 1:**
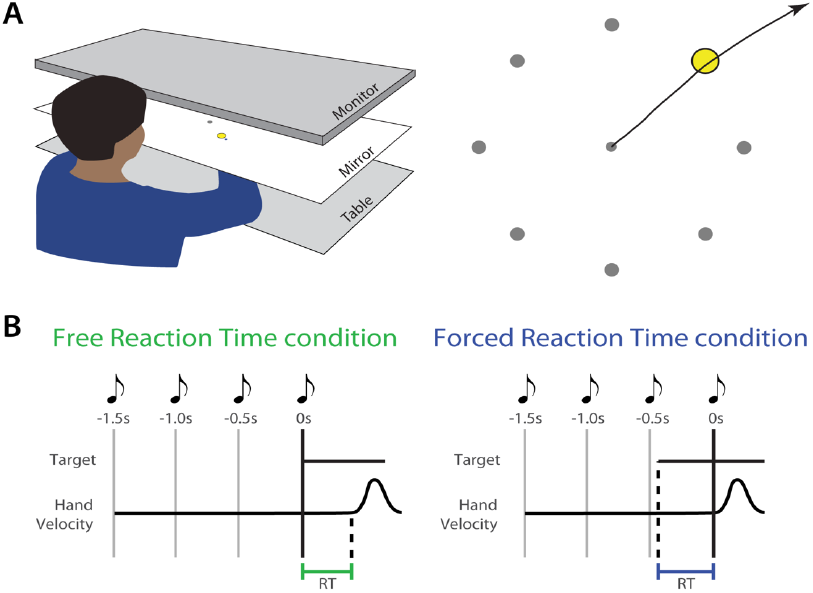
Apparatus and Experimental Conditions. A) Participants made planar reaching movements to interact with an on-screen display. Participants made ballistic ‘shooting’ actions with the goal of passing the cursor through a target. The target appeared in one of eight locations. B) Experimental conditions. In the Free Reaction Time condition the target appeared at a fixed time cued by a sequence of tones. Participants attempted to respond by initiating a movement as soon as possible. In the Forced Reaction Time condition participants always initiated movements at a fixed time (synchronously with the final tone in a sequence of four). The target appeared at a random time prior to movement; the time between target presentation and the fourth tone therefore imposed a limited response time.

Participants moved their hand to control the position of a cursor (blue circle, 5mm diameter). Each trial began with the cursor in a central start position (green circle, 10mm diameter). The two experimental conditions (Free and Forced Reaction Time - see below) required participants to make a ballistic arm movement (i.e. movements that use feedforward control with little opportunity to make online corrections to their movement; (Hardwick et al. 2013) with the goal to pass the cursor through a target (grey circle, 25mm diameter). The target could appear in one of eight locations, each spaced equally around the start position at a distance of 80mm.

### Free Reaction Time Condition

Participants were instructed to react as quickly as possible to the appearance of a target. The timing of stimulus presentation was predictable, occurring synchronously with the final tone in a sequence of four equally spaced tones (500ms separation). This cuing reduced ambiguity regarding the timing of stimulus presentation, which reduces reaction times and their variability (Frith and Done 1986). Participants completed 1-4 blocks (each 96 trials) of Free Reaction Time trials (the number of blocks varied depending on the time available to test the participant). The targets appeared in a pseudorandom sequence, with each target appearing 12 times per block.

### Forced Reaction Time Condition

The Forced Reaction Time condition used an established paradigm that requires participants to respond at a prescribed time within each trial (Ghez et al. 1997; Haith et al. 2015a, 2015b, 2016; Hardwick et al. 2019). Participants heard a sequence of four equally spaced tones (500ms separation), and were trained to initiate their movements synchronously with the onset of the fourth and final tone. Different reaction times were imposed by varying the time at which the target was presented relative to the required time of movement onset. Participants were instructed that while both the timing and the accuracy of their movements was important in this condition, their highest priority was to attempt to begin their response synchronously with the fourth tone. If participants failed to initiate their movement within +/−75ms of this time, on-screen feedback informed them that they were “Too early” or “Too late”. If participants failed to time their movement accurately on three consecutive trials the experimenter also provided additional feedback, reiterating the instruction that accurate timing was their highest priority in this condition. Analyses accounted for discrepancies in participant timing (i.e. differences in time between participants responses and the fourth tone) in several ways. First, we determined the ‘actual’ time participants used in each trial by measuring the time between the onset of the stimulus and their response (rather than the experimentally ‘prescribed’ time based on the time between stimulus onset and the fourth tone). Secondly, a set of ‘asynchrony’ analyses examined differences in timing between the participant responses and the fourth tone.

In initial training blocks the target appeared at the onset of the trial, allowing the participant 1500ms to prepare a response. Participants trained for one block of 50 trials; if they could accurately time the initiation of their movement in at least 35/50 trials they proceeded to the main experiment, otherwise they completed a second 50-trial training block. Participants then completed trials with variable target presentation times. In each block, target presentation varied uniformly between 0 and 400ms prior to the fourth tone (if participants failed to produce correct responses within this time window the range was increased to 600ms). Each block began with two ‘warm up’ trials in which the target appeared with the first tone. Participants completed 2-4 blocks (106 trials each) of Forced Reaction Time trials (the variable number of blocks depended on the time available to test the participant and their adherence to instructions).

### Data Analysis

Hand position was processed with a second order Savitzky-Golay filter (half-width 54ms). Movement onset was calculated as the time at which tangential hand velocity first exceeded 0.02m/s. We subtracted the mean delay in the recording system (measured to be 100ms) to provide a more accurate measure of true reaction time. Reaction time in both the Free Reaction Time and Forced Reaction Time conditions was calculated as the delay between the onset of the stimulus and movement onset. Initial movement direction was calculated from the direction of the hand’s velocity 100ms after movement onset.

Data from the Forced Reaction Time condition was used to model the probability of initiating an accurate movement at a given reaction time (i.e. a speed-accuracy trade-off) based on a previously established approach (Haith et al. 2016; Hardwick et al. 2019). Movements were considered to have been initiated in the correct direction if the initial movement direction was within 22.5° of the target. For data visualization purposes, the proportion of movements initiated in the correct direction was calculated for a 20ms sliding window around each potential reaction time. For analysis, a speed-accuracy trade-off was modeled as a cumulative Gaussian distribution centered on time T_p_ (thus T_p_ ~ N(U_p_, σ_p_^2^). This assumes movements before T_p_ were directed randomly with respect to the true target location, while movements after T_p_ were initiated in the correct direction with some probability α. Parameters were estimated from the data for each individual participant using a maximum likelihood approach.

Statistical analyses were conducted using JASP (0.13.1.0). The relationship between movement preparation and initiation was analyzed using a repeated-measures ANOVA (_RM_ANOVA). The _RM_ANOVA assessed the within-subjects factor of Time - Initiation Time (calculated using the Free Reaction Time condition) was compared to Preparation Time (calculated using the Forced Reaction Time condition), with Age included as a covariate. Further correlation and regression analyses assessed whether Age affected Initiation Time, Preparation Time, or the delay between them (i.e. Initiation Time minus Preparation Time). Data submitted to correlation analyses were screened for outliers using the “Robust Correlation” MATLAB toolbox (Pernet et al. 2013). This toolbox provides an objective approach to identifying and removing outliers without loss of statistical power. Where outliers were identified we report the ‘Skipped’ Pearson correlation (calculated by removing outliers and determining the correlation for the remaining datapoints), which directly reflects Pearson’s *r* (Pernet et al. 2013). Note that the inclusion/removal of outliers did not change any of our empirical results. Where appropriate, additional Bayesian analyses were conducted to determine the level of evidence in support of the null hypothesis (BF_01_), with classifications according to Wagenmakers et al. 2011. In the Bayesian analyses, outliers were removed based on the Robust Correlation procedure outlined above. Again, the inclusion/removal of outliers did not change any of our empirical results.

A series of control analyses examined the effects of the different experimental conditions, and participant age, on behavior. We first conducted correlation and regression analyses to determine whether participants completed the Free and Forced Reaction Time conditions with similar peak movement velocities. Possible differences were considered in a _RM_ANOVA comparing peak movement velocity across conditions (Free vs Forced Reaction Time conditions), including Age as a covariate. Additional correlation and regression analyses considered the relationship between participant Age and peak movement velocity in the Free and Forced Reaction Time conditions. Further analyses examined possible effects of Age on participant behavior in the Forced Reaction Time condition. Possible effects of Age on asymptotic accuracy (identified based on the model fit to the data for each participant) were examined using correlation and regression analyses. Possible effects of Age on timing accuracy were also assessed; Response Asynchrony was calculated as the difference in time between the fourth tone and the start of the participant’s response (Vleugels et al. 2020). Negative values therefore corresponded to moving before the fourth tone, and positive values corresponded to moving after the fourth tone. Correlation and regression analyses then assessed the possible relationship between Age and both signed and absolute Response Asynchrony.

All regression analyses are presented with bootstrapped 95% confidence intervals, calculated using resampling with replacement (Hardwick and Celnik 2014). A linear model was fit to each resampled population, and a line of best fit was then interpolated from the model parameters. This process was repeated 10,000 times, with the 2.5 and 97.5 percentiles of the interpolated fits being used as confidence intervals.

## Results

### Initiation time and preparation time dissociate

In line with our previous work, we found a significant difference between Initiation Time, as measured using the Free Reaction Time condition, and Preparation Time, as measured using the Forced Reaction Time condition, F_1,52_=77.7, p<0.001 (see Figure 2 for example data). Participants’ reaction times were significantly longer than the time they needed to prepare an accurate action in the Forced Reaction Time condition (t=24.82, p<0.01, mean Initiation Time (Free Reaction Time condition) = 290±34ms, mean Preparation Time (Forced Reaction Time condition) = 195±26ms, mean difference = 94±28ms).

**Figure 2:**
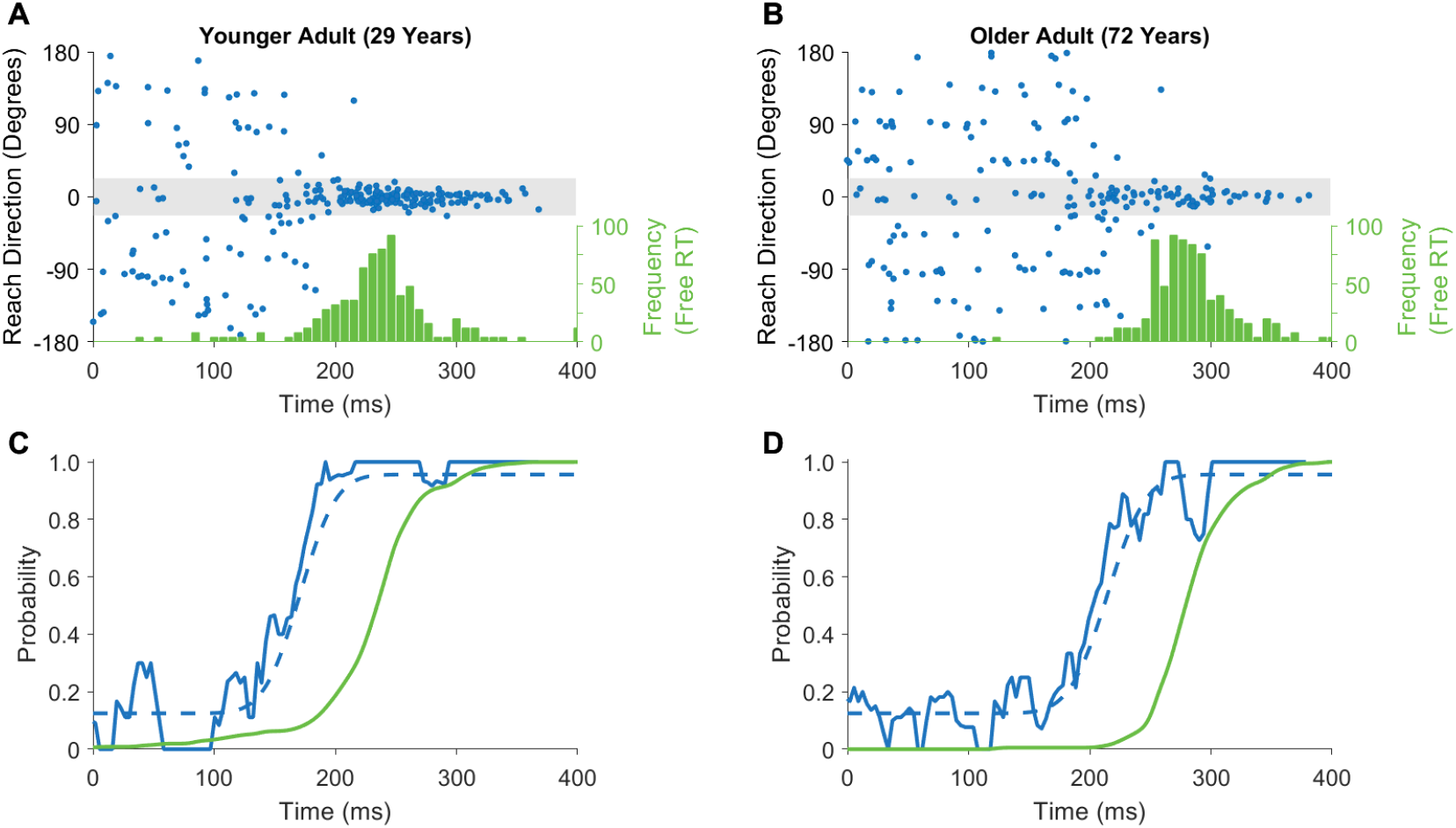
Data from example participants. Upper panels (A-B) show the distribution of reaction times in the Free Reaction Time condition (green histogram) and responses from individual trials in the Forced Reaction Time condition (blue dots). Responses falling within the grey shaded area were initiated in the correct direction. Lower panels (C-D) show a processed version of the data for the subject above. The solid green lines present a cumulative distribution of reaction times from the Free Reaction Time condition. Blue lines present data from the Forced Reaction Time condition; solid blue lines show a sliding window of successful responses, while dashed blue lines represent model fit to the data based on a cumulative Gaussian.

### Both initiation time and preparation time increase with age

While Age was not a significant covariate in the _RM_ANOVA for within-subject comparisons of Reaction and Preparation Times (F _*1,52*_=0.032, p=0.86), between-participants comparisons indicated that response times increased significantly with Age (F_*1,52*_ =8.0, p=0.007). Further analyses assessed the correlation between Age, Reaction Time, and Preparation Time. Replicating the findings of previous research, we found that increased age was related to a significant increase in reaction times in the Free Reaction Time condition (1 outlier removed, Skipped Pearson’s *r*=0.30, p=0.03; Figure 3A). Analysis of data from the Forced Reaction Time condition also revealed that movement preparation time increased significantly with Age (2 outliers removed, Skipped Pearson’s r=0.45, p=0.0007; Figure 3B). Accuracy in the Free Response condition high for all participants (mean 98.9±1.3%), and analysis indicated there was no significant correlation between accuracy and age (r=−0.08, p=0.56). Further Bayesian correlation analysis found substantial evidence for the null hypothesis (BF_01_=5.0), indicating that performance in the Free Response condition was close to ceiling for all participants, regardless of their age.

**Figure 3:**
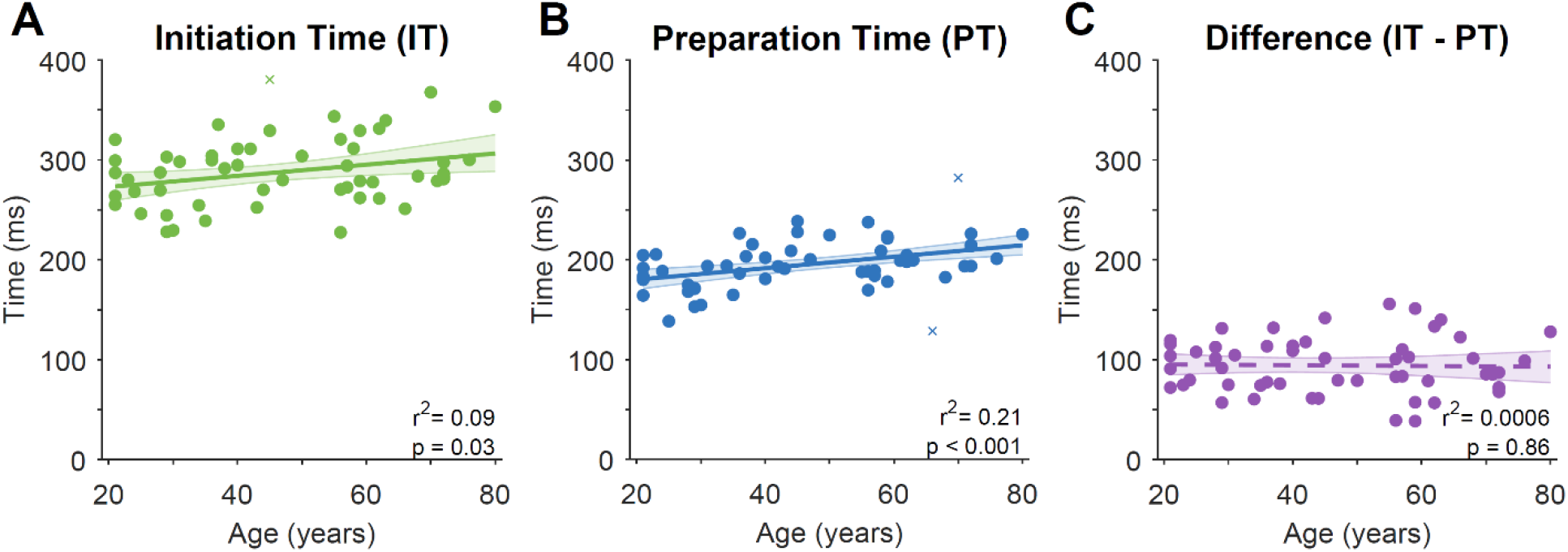
Relationships between Age and movement Initiation Time (Free Reaction Time condition), Preparation Time (Forced Reaction Time condition), and the delay between movement Preparation and Initiation. Each point presents data from a single subject (crosses indicate outliers as identified by robust correlation analysis, which were not included in summary statistics). Solid line presents linear regression on the data, dashed lines present non-significant regression lines. Error bars present bootstrapped 95% confidence intervals.Age does not affect the delay between movement preparation and initiation

### Age does not affect the delay between movement preparation and initiation

The delay between movement preparation and initiation was calculated for each participant by taking their mean reaction time, as established in the Free Reaction Time condition, and subtracting their mean preparation time, established based on the speed-accuracy trade-off observed in the Forced Reaction Time condition (Figure 4). As identified in an earlier analysis, all participants exhibited a delay between movement Preparation and Initiation (mean±SD = 94±28ms). There was, however, no significant relationship between age and the duration of the delay (Figure 3C, Pearson’s *r*=−0.025, p=0.86). Further analysis using Bayesian correlation indicated there was substantial support for the null hypothesis (BF_01_ = 5.801)(Wagenmakers et al. 2011).

**Figure 4:**
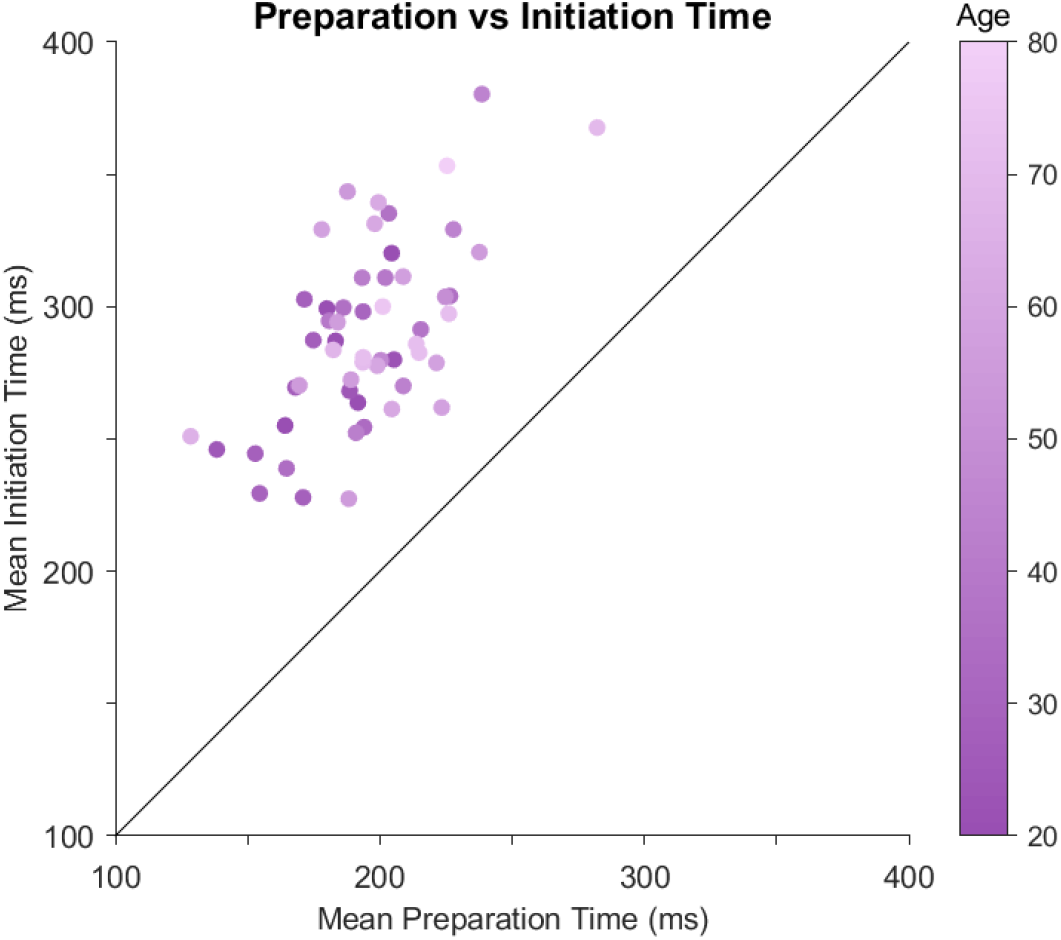
Preparation Time vs Initiation Time. Each circle represents one participant, with lighter colors presenting increasingly older participants. Note that each participant’s Initiation Time (average of reaction times for that participant in the Free Reaction Time condition) was greater than their Preparation Time (average time of response preparation based on a model fit to data for that participant in the Forced Reaction Time condition).

### Peak movement velocity was correlated across conditions and decreased with age

Control analyses examined whether peak movement velocity affected performance within and across conditions. Participant peak movement velocity in the Free and Forced Reaction Time conditions was highly correlated (8 outliers removed, Skipped Peason’s *r* = 0.79, p=5.7916e-11 Figure 5A). A corresponding _RM_ANOVA found no significant difference between peak movement velocity in the Free and Forced Reaction Time conditions (_RM_ANOVA, F_*1,52*_ =0.87, p=0.36), suggesting participant movement speeds were consistent between the two conditions. As older age is associated with slower movement speeds, we also examined whether peak movement velocity differed with Age. Age was not a significant covariate in the _RM_ANOVA (F_*1,52*,_ =0.31, p=0.58), but the analysis indicated a trend for Age as a between-subjects effect on peak velocity (_RM_ANOVA, F_*1,52*_=3.7, p=0.06). Correlation analyses suggested that peak velocities increased with age, with trends for this effect in both the Free Reaction Time condition (Pearson’s *r*=−0.26, p=0.055; Figure 5B) and Forced Reaction Time condition (Pearson’s *r*=−0.24, p=0.088; Figure 5C).

**Figure 5:**
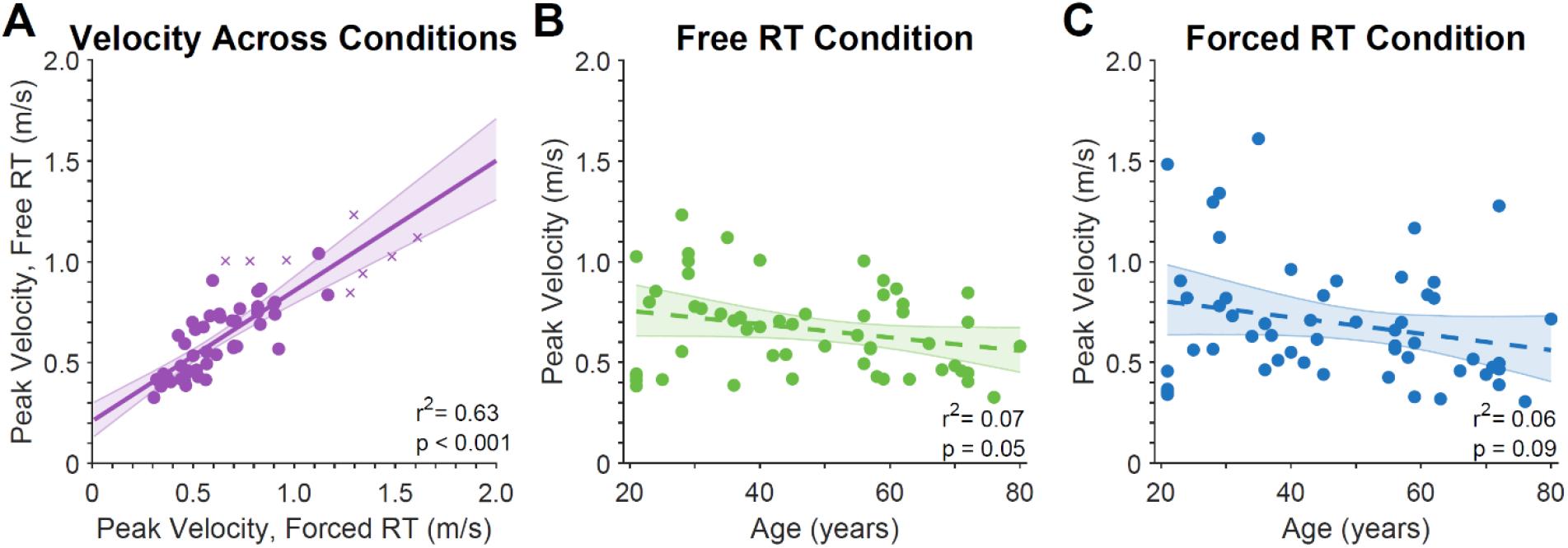
Analyses of peak velocity. Left panel shows correlation between Peak velocity in the Free and Forced Reaction Time conditions. Central and Right panels show correlations between peak velocity and age in the Free and Forced Reaction Time conditions, respectively. Each point presents data from a single subject (crosses indicate outliers as identified by robust correlation analysis, which were not included in summary statistics). Solid line presents linear regression on the data, dashed lines present non-significant regression lines. Error bars present bootstrapped 95% confidence intervals.

Further analysis examined whether differences in movement speed across ages might have accounted for the observed differences in preparation time and initiation time. We found no significant relationship between reaction time and peak velocity in the Free Reaction Time Condition (Pearson’s *r*=−0.14, p=0.30; Figure 6A), or the Forced Reaction Time Condition (1 outlier removed, Skipped Pearson’s *r*=−0.18, p=0.19, Figure 6B). Bayesian analysis indicated there was substantial support for the null hypothesis when comparing reaction time and peak velocity in the Free Reaction Time condition (BF_01_=3.5), and anecdotal evidence for the null hypothesis in the Forced Reaction Time condition (BF_01_=2.6).

**Figure 6:**
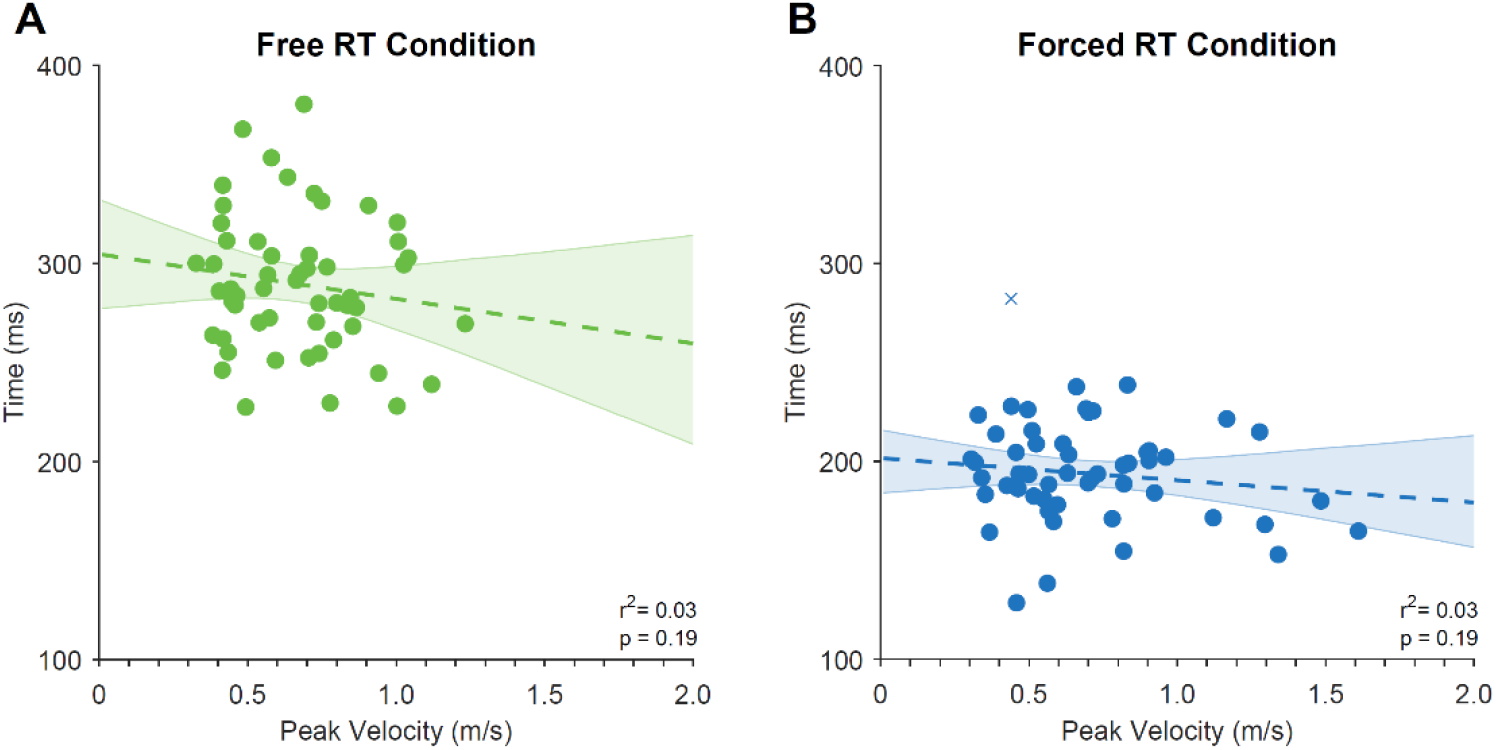
Comparisons of peak velocity and reaction time for the Free Reaction Time condition (A) and Forced Reaction Time (B) conditions. Each point presents data from a single subject (crosses indicate outliers as identified by robust correlation analysis, which were not included in summary statistics). Dashed lines present non-significant regression lines. Error bars present bootstrapped 95% confidence intervals.

### Asymptotic accuracy in the Forced Reaction Time condition decreased with age

A correlation analysis indicated that asymptotic accuracy in the Free Reaction Time condition decreased significantly with age (r=−0.34, p=0.012; see Figure 7A). This decline occurred at a relatively low rate (0.0017% decrease in accuracy per year), corresponding to an approximate decrease of 11% from ages 20 to 80 (97% vs 86% accuracy, respectively).

**Figure 7:**
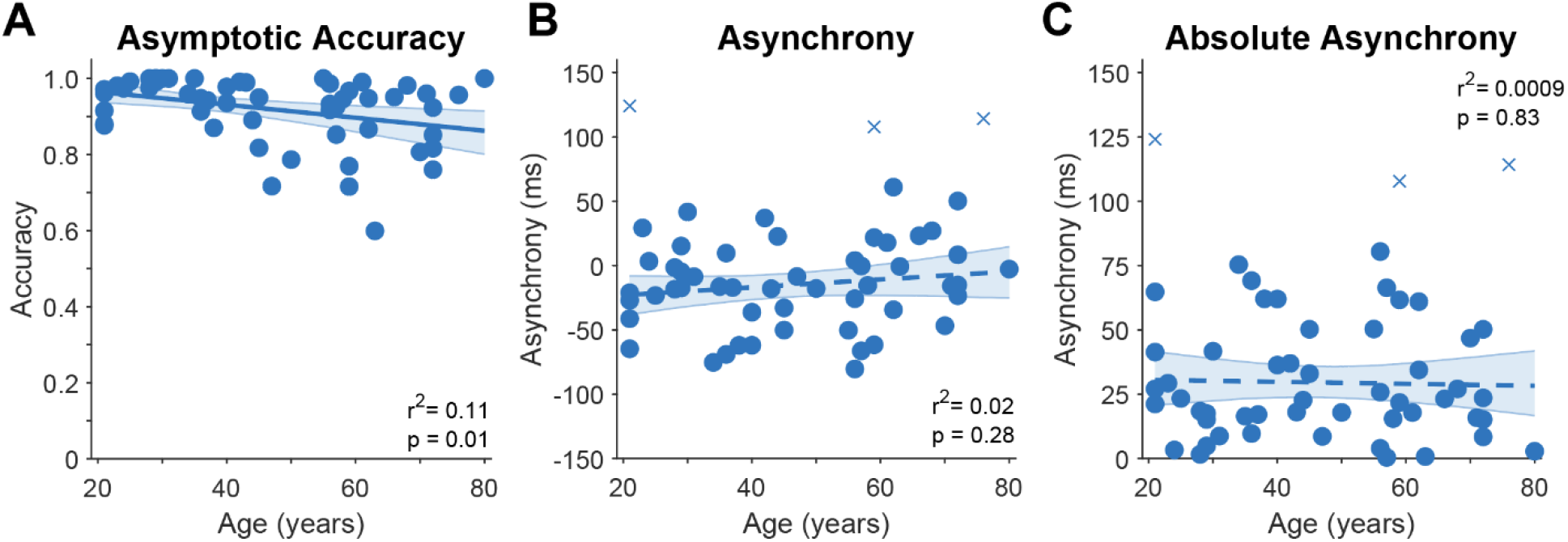
Effects of Age on behavior in the Forced Reaction Time condition. Left panel indicates the significant relationship between Age and Asymptotic Accuracy. Right panel indicates the non-significant relationship between Age and Response Asynchrony. Each point presents data from a single subject (crosses indicate outliers as identified by robust correlation analysis, which were not included in summary statistics). Solid line presents linear regression on the data, dashed lines present non-significant regression lines. Error bars present bootstrapped 95% confidence intervals.

### Timing (Asynchrony) in the Forced Reaction Time condition did not differ with age

A final analysis examined participant’s ability to time their responses in the Forced Reaction Time condition to coincide with the fourth tone. Signed response Asynchrony did not differ significantly with age (Pearson’s *r*=0.15, p=0.29, 3 outliers removed, Skipped Pearson’s *r*=0.16, p=0.28: See Figure 7B), and Bayesian analysis provided substantial evidence in support of the null hypothesis (BF_01_=3.2; Wagenmakers et al. 2011). Absolute response Asynchrony also did not differ with age (Skipped Pearson’s *r*=−0.03, p=0.83), with further Bayesian analysis again providing substantial support for the null hypothesis (BF_01_=5.6). Together these analyses suggest that timing asynchrony in the forced response condition did not differ significantly with age.

## Discussion

We used a visually-guided planar reaching task to measure reaction times and assess the time participants needed to prepare accurate movements. In line with previous studies, we found that ‘Free’ reaction times increased linearly with age (Fozard et al. 1994; Gottsdanker 1982; Woods et al. 2015). We compared these data to performance in a ‘Forced Reaction Time’ condition, in which we measured the minimum time participants required to prepare accurate movements by forcing them to respond with shorter-than-normal response times. The time required to prepare accurate movements also increased linearly with age, and was significantly shorter than the reaction time, replicating our previous observation that movements are not immediately initiated once they are prepared (Haith et al. 2016). Further analysis identified that age had no significant effect on the delay between movement preparation and initiation. These results indicate that the slower reaction times of healthy older adults observed in this task were not due to an increased hesitancy to respond, but can instead be wholly attributed to declines in the ability to process stimuli and prepare accurate movements.

Healthy human aging is associated with changes in motor behavior including declines in coordination, increased kinematic variability, and a reduced ability to modify movements to respond to changes in the environment (Hardwick and Celnik 2014; Sarlegna 2006). Such age-related changes in behavior are accompanied by changes in brain structure and function (Dully et al. 2018; Heuninckx et al. 2008; Stadlbauer et al. 2008). The increase in the amount of time required to prepare movements with age, as identified here, is consistent with these previous findings. Previous work has also suggested that healthy older adults prefer to respond with longer reaction times to ensure accurate responses (Salthouse 1979; Smith and Brewer 1995; Starns and Ratcliff 2010). Here we found no evidence of such age-related delays in responding. We note, however, that the simple reaching task used here had relatively low cognitive demands. Age-related declines in performance are exacerbated by increased task complexity and/or greater cognitive demand (Woods et al. 2015), consistent with frequently demonstrated differences between cognitive and motor functions (Wollenweber et al. 2014; Wu et al. 2004). We therefore propose that the reported delaying of action in those studies may not represent a ‘default policy’ for older adults, but could instead occur in response to increases in task complexity.

Further analyses indicated that increasing age was associated with slower peak movement velocities in all conditions, and decreases in asymptotic accuracy in the Forced Reaction Time condition. This drop in accuracy may have reflected an increased propensity for lapses in concentration, particularly given the dual demands of timing and accuracy in the Forced Reaction Time condition. Skilled motor performance is characterized by both speed and accuracy (Hardwick et al. 2017; Rajan et al. 2019; Reis et al. 2009; Shmuelof et al. 2012, 2014), and the present data are consistent with aforementioned and well-established age-related declines in movement control. By contrast, there was no significant effect of age on the ability to synchronize responses with the fourth tone, as evidenced by the analysis of Response Asynchrony in the Forced Reaction Time condition. Note, however, that this does not necessarily reflect spontaneous, self-selected participant behavior. Instructions to participants in the Forced Reaction Time condition emphasized that while both the accuracy and timing of their responses were important, timing was the highest priority. Older adults may have had greater asynchrony (due to a tendency to delay their movements to wait for the target to appear, so they could reach in the correct direction) without this intervention. We therefore conclude that increasing age was associated with a decrease in overall performance (i.e. older adults had longer Initiation Times, longer Preparation Times, lower peak movement velocities, and were less accurate).

In summary, our results are consistent with previous observations that humans delay the initiation of prepared movements, and show that the size of this delay remains constant across the lifespan. The consistent duration of this delay indicates that healthy older adults do not appear to change their behavior in relatively simplistic response time tasks in order to favor accuracy at the expense of speed. The declines in their performance observed here can instead be wholly attributed to age-related changes in their capability to process and prepare movements.

## Acknowledgements

This work was supported by NSF grant 1358756. This project has received funding from the European Union’s Horizon 2020 research and innovation programme under the Marie Skłodowska-Curie grant agreement No 702784 (RMH). RMH is supported by grants from the UC Louvain special research fund (1C.21300.057 and 1C.21300.058). We thank Jeff Gooding for assistance with data collection.

## Author Contributions

RMH conceived the research. RMH, AF, and MGC collected the data. RMH analyzed the data.

RMH drafted the manuscript. RMH, AF, MGC, KZ and AH revised the draft.

## Additional Information

The authors declare no competing interests.

## Data Availability

The datasets generated during and/or analyzed during the current study are available from the corresponding author on reasonable request.

